# Disturbance increases functional diversity but decreases phylogenetic diversity of an arboreal tropical ant community

**DOI:** 10.1101/2023.04.13.536723

**Authors:** Philipp O. Hoenle, Nichola S. Plowman, Pável Matos-Maraví, Francesco de Bello, Tom R. Bishop, Martin Libra, Cliffson Idigel, Maling Rimandai, Petr Klimes

## Abstract

1. Tropical rainforest canopies host a highly diverse arthropod fauna, which contribute to ecosystem function through their functional (FD) and phylogenetic diversity (PD). While a lot of previous research has documented the severe negative impacts of disturbance on the FD and PD of ground invertebrate communities, our understanding of arboreal counterparts is limited.
2. Here, we studied the effects of forest disturbance on an ecologically important invertebrate group, the ants, in a lowland rainforest in New Guinea. We exhaustively sampled 4000 m^2^ area of a primary and a secondary forest for canopy ants. We report > 2800 occurrences of 128 ant species in 852 trees, one of the most comprehensive arboreal collections to date.
3. To test how ant PD and FD differ between the two forests, we constructed the ant species-level community phylogeny and measured 10 functional traits. Furthermore, we assessed by data exclusion the influence of species which were not nesting in individual trees (visitors) or only nesting (nesters), and of non-native species on FD and PD values. We expected that disturbance would decrease FD and PD in tree dwelling ants. We hypothesized that traits in primary forests would be more overdispersed due to the greater availability of ecological niches, while secondary forests would have stronger trait clustering due to a a stronger habitat filtering caused by more extreme microclimate.
4. Primary forests had higher species richness and PD than secondary forest. Surprisingly, we found higher FD in secondary forest. This pattern was robust even if we decoupled functional and phylogenetic signals or if non-native ant species were excluded from the data. Visitors did not contribute strongly to FD, but they increased PD. Community trait means further corroborate the functional distinctiveness of arboreal ants among secondary and primary forest, with almost all traits being impacted by disturbance and forest succession.
5. We conclude that the most plausible explanation is increased competition among closely related ant species in the secondary forest, which drives trait divergence. In the primary forest, abiotic habitat filters leads to more similar morphology and thus lower FD of phylogenetically more diverse ant assemblages.

## Introduction

Understanding the consequences of disturbance on biological diversity remains a challenge (Whittaker et al., 2001; Naeem et al., 2012; Ewers et al., 2015), particularly in the face of the global anthropogenic landscape transformation (Hansen et al., 2013). While the negative effects of human disturbance on taxonomic diversity (TD) are well documented (Lawton et al., 1998; Dunn, 2004), their consequences for ecosystem functioning and community assembly are much less understood.

A promising way to measure disturbance effects on ecosystems and their biota is the study of community traits and phylogenies between habitats that were exposed to different levels of disturbance. Functional traits link the performance of species to their environment, and the diversity of these traits (functional diversity, FD) is therefore assumed to reflect ecosystem function (Diaz et al., 2001; Cadotte et al., 2010). In addition, phylogenetic diversity (PD) captures the evolutionary history of a community and therefore provides complementary insights on the consequences of disturbance (Flynn et al., 2011; Srivastava et al., 2012; Tucker et al., 2018). Measures of FD and PD are interdependent and often positively correlated (Cadotte et al. 2019). For instance, in a neutral scenario of community assembly, species in a phylogenetically more diverse assemblage are accumulating greater variation in the ecologically important traits as opposed to a phylogenetically more closely related assemblage (Webb et al., 2002; Prinzing et al., 2008).

In the majority of studies, FD and PD increase with rainforest succession, suggesting a decrease in ecosystem function under human disturbance (Whitfeld et al., 2012; Bu et al., 2014; Letcher et al., 2012; Mo et al., 2013). A similar pattern holds for TD (Dunn, 2004), although its decline with disturbance is usually more pronounced compared to PD and FD (Liu et al., 2019, Salaz-Lopez, 2017). However, decreases in diversity measures with disturbance are not universal and taxa can show idiosyncratic responses. For instance, among lowland tropical birds, Sreekar et al. (2021) found that FD increased through disturbance, while PD remained stable.

Comparing both FD and PD after disturbance can provide insights into underlying community assembly processes (Cavender-Bares et al., 2009). For instance, if a community is clustered in its traits, it potentially indicates niche-based habitat filtering, since only species with traits suitable to the environment are selected and thus more uniform than expected at random (Kraft et al., 2015). Overdispersion, on the other hand, can be an indicator for competitive exclusion, where communities are more dissimilar in their traits in order to mitigate competition (limiting similarity principle; MacArthur and Levins, 1967).

In tropical forests, vertical stratification drives insect diversity and abundance (Xing et al., 2022). For instance, the canopy ant fauna makes up about half of the total diversity and a significant contribution to the invertebrate biomass (Floren et al., 2014; Davidson et al., 2003). Ants display distinct lifestyles and morphology fitted to a life in trees (Basset et al., 2015; Leponce et al., 2021) and play multiple ecological roles as detritivores, predators and herbivores (Lach et al., 2010). Yet, unlike ground-dwelling ants (e.g., Bihn et al., 2010; Liu et al., 2016), we know little about how disturbance shapes the functional and phylogenetic structure of arboreal communities. Arboreal ant communities consist of the ants that nest on a focal tree, and also of ants that come to search for food but nest elsewhere (either on the ground or the surrounding vegetation). Through altering, for instance, the nesting resources and connectivity, disturbance might change the contribution of these two components of the tree. As a result, the fraction of species coming from ground level versus true arboreal nesters may change with disturbance, and thus alters their contribution to tree scale functional and phylogenetic diversity.

Furthermore, disturbance is associated with an influx of non-native species. While there is only little previous research on the functional consequences of ant invasion (but see Wong et al., 2020), it is well established that non-native species can have devastating consequences for ecosystem function (Lach et al., 2010) and lower PD and FD of communities (Loiola et al., 2017). Moreover, canopy specialists are expected to be more sensitive to structural and climatic changes than ground-dwelling species (Klimes, 2017; Parr & Bishop, 2022). Thus, there is an uncertainty around the consequences of forest disturbance on a key ecological group.

Here, we report on the first investigation of the impact of human disturbance on the PD and FD of arboreal ants, using a diverse tropical lowland forest from Papua New Guinea as study system. We already know from this particular forest that human disturbance lead to drastic species composition changes and taxonomic diversity declines (Klimes et al., 2012, 2015). However, the consequences of these changes on the PD and FD of tree-dwellig ants are unclear. We constructed a species-level community phylogeny and assessed their morphological variation in functional traits to answer this question. Compared to previous studies on arboreal ants which sampled either a few trees or individual branches and epiphytes (e.g., Fayle et al., 2013: Floren et al., 2014), our approach represents a complete census of 852 whole tree communities within 4.000 m^2^ of a primary and a secondary (human-disturbed) forest (Klimes et al., 2012, 2015). The data was collected destructively by felling and dissecting trees in the plots targeted for a slash-and-burn agriculture in collaboration with local landowners while supporting conservation of the site (Novotny, 2010).

We predict that arboreal ants in secondary forest will have lower FD and PD than in primary forest. Vegetation structure and nesting opportunities for ants in the secondary forest are less complex (Klimes, 2017; Mottl et al., 2019), and the microclimate tends to be warmer and more variable (Jucker et al., 2018). As a result, we expect that ant morphology will reflect this reduction in niche diversity and stronger microclimatic filters, leading to higher clustering in traits and phylogeny in secondary forest communities. In contrast, we anticipate neutral or competitive processes (i.e., overdispersion in traits and phylogeny) to be prevalent in primary forest due to spatial exclusion (i.e., competition) by the most common canopy species of these primary forests (Leponce, et al. 2021; Mottl et al. 2021). Furthermore, habitat filtering and competitive interactions are expected to lead to differences in several key ant functional traits between both forest types (Martello et al., 2018). For instance, undisturbed habitats are known to support a wide range of ant body sizes (Gibb et al., 2017b). Further, we expect that species that do not nest in the sampled trees but forage on them for food resources and thus boost those trees’ ant richness (see Klimes et al. 2015) will substantially contribute to increased functional and phylogenetic diversity in both forest types. Finally, we expect that one of the main drivers of lower functional and phylogenetic diversity will be non-native ant species that invaded the secondary forest.

## Materials and Methods

### Study site

Our study was based in a lowland rainforest near Wanang Conservation Area, Madang Province, Papua New Guinea (100 – 200 m above sea level; 05 ° 14’S 145 ° 11’E). The region experiences a mean annual rainfall of 3600mm, a mean annual temperature of 26.5⁰C and a weak dry season from July to September (McAlpine et al., 1983). As part of a larger study, 1-ha each of primary forest and secondary forest were felled in collaboration with local landowners (see Klimes et al., 2012; Whitfeld et al., 2012; Whitfeld et al. 2014). We exhaustively sampled 0.4ha (100 x 40 m) within each hectare plot for arboreal ants; in primary forest (an old growth forest stand of minimal 50 years without human disturbance), and in secondary forest with *∼*10 years of secondary regrowth after the abandonment of small-scale slash-and-burn agriculture. Our dataset is thus a spatially extended version of the data used in previously published studies on taxonomic diversity of ants from these felled plots (0.3 ha with dimensions 80 x 40 m in each forest; Klimes et al., 2012, 2015).

In the two plots, all trees with diameter above breast height (DBH) ≥ 5cm were felled, measured, and identified to species. We exhaustively sampled the arboreal ant communities by inspecting trees immediately after felling. From the tree base to the tree crown, we searched each tree for ants, dissecting bark, branches, trapped soil and litter and epiphytes to find the more cryptically nesting and foraging individuals. In each tree, we recorded whether ant species were collected from nests or were foraging from elsewhere (i.e. nesters and visitors – see below for definition). A full sampling protocol for vegetation and ants is available in Volf et al. (2019) and the details on vegetation characteristic of the two plots are available in Whitfeld et al. (2012). We hand collected ants of all castes available and representative specimens were stored in ethanol along with tree and nest identifiers. Only established nests with workers and worker foragers were considered in the analysis in this study. Singleton queens, and those with brood but without workers, were excluded as they did not have established colonies, and thus unlikely to have an important ecological impact in the community, and are thus not considered for trait measures (Parr et al., 2017).

### Ant identification and phylogeny

Ant individuals were sorted to genus and to species or morphospecies (species hereafter) using available taxonomic keys (Bolton, 1994; Andersen, 2000; Schmidt & Shattuck, 2014), collections at the Institute of Entomology (Biology Centre, Czech Academy of Science), the Museum of Comparative Zoology (MCZ) at Harvard University, online databases (http://www.antweb.org), and the assistance of specialist taxonomists (see acknowledgements). For each species, representative workers were sequenced for the mitochondrial gene fragment cytochrome c oxidase I (COI, 659 bp) and the nuclear gene fragment *wingless* (Wg, 409 bp). We obtained COI and Wg sequences for 112 of 127 species, we gathered missing COI and/or Wg sequences for further 4 species from the Genbank database, while the remaining 11 species were represented by a single gene (6 COI, 5 Wg; 8% of species). Details on the extraction process and phylogenetic tree assembly can be found in the Supplementary Text S1.

### Trait measurement

Most studies of ant functional diversity are based only on the traits of minor workers, because the most common sampling methods do not reliably sample all castes of ant species. However, intraspecific variability of ants is important to consider (Wong and Carmona, 2021), and our whole forest destructive sampling method allowed us to collect all distinct size classes of almost all ant species present in the nests and foragers. Thus, we measured all available worker and soldier castes of 1-11 point-mounted individuals of each species (581 individuals, mean 4.5 individuals per species).

We followed the guidance in previous studies to choose ecologically relevant traits and their standard measurements (Gibb et al., 2015; Parr et al., 2017). We measured the following continuous traits: head length, head width, clypeus length, leg length (=hind femur + tibia length), eye size (approximated as area of an ellipse using eye length and width), eye position (=head width - interocular distance), spinosity (total number of spines on alitrunk and petiole), and mandible length. Traits were measured to the nearest 0.01mm using an Olympus SZX7 stereomicroscope with magnification 18x to 126x, fitted with an ocular graticule. For each species we further recorded a categorical trait, the degree of cuticular sculpturing (0=none, 1=shallow, 2=moderate, 3=deep). Finally, we introduce a new ant trait which considers the degree of polymorphism, which is calculated as the maximum head length (a proxy for body size) of a species divided by the minimum head length (among all castes). Thus, species with no intraspecific variation have polymorphism values close to 1, while polymorphic species can have a value of up to 2.6 (e.g., in *Pheidole* species).

Since all continuous measurement traits are highly correlated with body size, we calculated relative measures using head length as an appropriate surrogate for body size (Gibb et al., 2017b; for size correated correlations, see Figure S1.). We then calculated the mean value of each trait for each species. To account for differences in the relative number of minor and major workers in strongly polymorphic species, we calculated a weighted mean for all size-related traits. We use a conservative weighing of a caste-ratio of 20% major and 80% minor workers (e.g., Tschinkel, 2005; Walker and Stamps, 1986). To measure overall functional diversity, including the multiple functional traits, we calculated the Gower distance of categorical and continuous traits combined, using the function *gowdis* in the R package FD (Laliberté et al., 2010).

### Statistical Analyses

We are aware that our design is pseudoreplicated since we are comparing two forest plots. This is a common problem for whole-ecosystem approaches with very abundant taxa such as ants, but arguably the careful use of inferential statistics is possible (Oksanen, 2001; Chaves, 2010). We know both forests plots possess vegetation structure and tree species composition typical for primary and secondary forest in our studied site (Whitfeld et al., 2014). Since we are investigating ants, we are interested in a comparative small scale, as most ant colonies forage in one or few trees at most (with the exception of few supercolonial species). Therefore, statistically treating tree-level communities as independent replicates poses a minor problem, particular considering our large plot size encompassing hundreds of trees and a high data quality through exhaustive sampling.

A tree harbours two distinct ant communities. There are ants which nest on the tree (’nesters’), and those which forage on the tree but either have their nest on neighbouring trees or on the not sampled shrubs and forest ground level (’visitors’, i.e., equal to the “F-N” foraging species in Klimes et al. 2015). Nesters use the trees both for nesting space and food resources, while visitors use only the latter (Klimes, 2017). Our analysis untangles the functional and phylogenetic contribution of each group by providing a separate analysis on nesters, visitors, and the whole community (’all’). We use three separate community matrices, which all contain ant species occurrences in each tree: ‘all’, ‘nesters’ and ‘visitors’. We calculated the visitor matrix community by subtracting the ‘nester’ matrix from the ‘all’ matrix. Hence, the ‘visitor’ matrix has no species overlap with the nesters, thus allowing to untangle the relative contribution of nesters and foragers to functional and phylogenetic structure of each tree. Note that an ant that nests on the tree does also forage there, but is not included in the ‘visitor’ matrix. A visitor ant that belongs to the same species as a nesting ant is also disregarded in the visitor matrix, since conceptually, it’s functional and phylogenetic contribution is already accounted for. Since the community matrices are derived from each other and thus violate assumptions of independence, we do not assess statistical differences between nesters, foragers, and the whole community within the same forest, but instead focus our statistical comparison on the differences between primary and secondary forest.

We analysed our data on two different scales, tree and plot scale. The tree scale analysis is based on the ant occurrences per tree, and thus gives tree-level values for each functional and phylogenetic diversity index. The plot scale analysis is based on the summed-up occurrences across all trees in each forest plot. The tree-scale analysis excludes communities with < 2 species, but in the plot-wise analysis all ant occurrences are retained. Most analyses were performed on both plot- and tree-scale, with the exception of the community weighted means (CWMs) of individual traits (only tree scale) and species composition overlap (only plot scale). We calculated species overlap on plot scale between nesters and foragers, as well as between primary and secondary forests using the Bray-Curtis distance using the package ‘vegan’ (Oksanen et al. 2020).

Based on either tree- or plot scale matrices, we calculated the Rao quadratic entropy (Rao Q) as indicator for PD and FD using the abundance-weighted function ‘ses.mpd’ from ‘picante’ (Kembel et al. 2010). Note that the abundance-weighted measure in this package calculates Rao Q when calculated on occurrence data (de Bello et al., 2016). For FD, we used a gower distance matrix of the functional traits. For PD, we used the square-root transformed phylogenetic distance tree of all ant species (see above).

To assess if our communities were phylogenetically clustered or overdispersed we calculated the standardised effect sizes (SES) and P values of observed Rao Q as compared to a null model based on taxa-swaps (shuffled distance matrix labels across all taxa included in the whole distance matrix; 999 runs). Negative SES values and low quantiles (p<0.05) indicate phylogenetic clustering, and positive SES values and high quantiles (p>0.95) indicate overdispersion (Table 1A; Götzenberger et al., 2016). Since the p-values obtained through the null-model comparison is regarded as conservative, we further test the distribution of SES against 0 with a Wilcoxon’s signed rank test for the tree-scale data (Table 1B). If the distribution is significantly lower than 0, we regard it as evidence for clustering, if it is significant higher than 0 we regard it as evidence for overdispersion. Note that observed values and SES obtained from the taxa-swap null model are highly correlated (de Bello, 2012; Götzenberger et al., 2016), which is also the case in our data. Therefore, for simplicity we interpret both FD and PD patterns and their assembly mechanisms through statistically comparing and plotting the SES values, while we provide the observed FD and PD values only in the Tables.

**Table 1.** Characteristics of arboreal ant communities sampled in 0.4 ha of primary and 0.4 ha of secondary lowland rainforest in Papua New Guinea, and of their taxonomic, functional (FD) and phylogenetic (PD) diversity. (A) values on plot scale and (B) values on tree scale (852 trees sampled in total). Positive Standardised effect sizes from null models (Rao Q SES) indicate community overdispersion (cells in light blue) and negative SES indicate clustering (cells in light orange). On plot scale, we give the p-values in comparison to the taxa-swap null model, while for the tree scale we test the distribution of all SES values against 0 with a Wilcoxon’s signed-rank test. Significant clustering and overdisersion values and their respective p-values are highlighted in bold (alpha = 0.05).

| **A) Plot scale** | **Primary all** | **Secondary all** | **Primary visitors** | **Secondary visitors** | **Primary nesters** | **Secondary nesters** |
| --- | --- | --- | --- | --- | --- | --- |
| N of sampled trees | 472 | 380 | 472 | 380 | 472 | 380 |
| N of species occurrences | 1678 | 1173 | 671 | 511 | 1007 | 662 |
| species richness | 101 | 57 | 64 | 34 | 84 | 50 |
| FD (Rao Q observed) | 0.191 | 0.212 | 0.188 | 0.209 | 0.207 | 0.206 |
| FD (Rao Q SES) | -1.042 | 0.035 | -0.819 | 0.212 | -1.067 | -0.414 |
| p value (null model) | 0.153 | 0.532 | 0.219 | 0.601 | 0.153 | 0.355 |
| decoupled FD (Rao Q SES) | -0.459 | 0.963 | -0.274 | 1.106 | -0.856 | 0.229 |
| p value (null model) | 0.398 | 0.821 | 0.491 | 0.846 | 0.203 | 0.625 |
| PD (Rao Q observed) | 0.531 | 0.518 | 0.515 | 0.483 | 0.539 | 0.520 |
| PD (Rao Q SES) | -0.025 | -0.665 | 0.297 | -0.714 | -0.690 | -0.063 |
| p value (null model) | 0.544 | 0.240 | 0.222 | 0.677 | 0.262 | 0.484 |
| decoupled PD (Rao Q SES) | -0.464 | -0.636 | -0.265 | -0.457 | -0.877 | -0.828 |
| p value (null model) | 0.314 | 0.272 | 0.400 | 0.318 | 0.184 | 0.213 |

| **B) Tree scale** | **Primary all** | **Secondary all** | **Primary visitors** | **Secondary visitors** | **Primary nesters** | **Secondary nesters** |
| --- | --- | --- | --- | --- | --- | --- |
| N of occupied trees | 442 | 345 | 362 | 263 | 324 | 276 |
| N of trees with $\geq 2$ species | 383 | 279 | 284 | 191 | 173 | 136 |
| mean species richness | 3.55 | 3.09 | 2.13 | 1.74 | 1.42 | 1.35 |
| mean FD (Rao Q observed) | 0.142 | 0.159 | 0.132 | 0.152 | 0.129 | 0.136 |
| mean FD (Rao Q SES) | **-0.498** | **0.061** | **-0.450** | 0.051 | **-0.391** | -0.051 |
| p value (against 0) | **< 0.001** | **0.047** | **< 0.001** | 0.259 | **< 0.001** | 0.755 |
| mean decoupled FD (Rao Q SES) | **-0.220** | **0.478** | **-0.096** | **0.562** | **-0.269** | 0.132 |
| p value (against 0) | **< 0.001** | **< 0.001** | **< 0.001** | **< 0.001** | **< 0.001** | 0.435 |
| mean PD (Rao Q observed) | 0.409 | 0.382 | 0.373 | 0.348 | 0.347 | 0.341 |
| mean PD (Rao Q SES) | **0.067** | **-0.367** | **0.169** | **-0.484** | -0.13 | 0.079 |
| p value (against 0) | **0.034** | **0.001** | **< 0.001** | **0.008** | 0.558 | 0.056 |
| mean decoupled PD (Rao Q SES) | **-0.129** | **-0.374** | -0.054 | -0.311 | **-0.164** | **-0.364** |
| p value (against 0) | **<0.001** | **< 0.001** | 0.382 | 0.15 | **0.012** | **< 0.001** |

PD and FD are closely related if traits are strongly phylogenetically conserved. To check this, we tested a measure of the phylogenetic signal of each trait by using Blombergs K (Blomberg et al., 2003). Further, we investigated the correlation between FD and PD on tree scale communities by using a linear model which includes forest type as predictor. Also, we investigated the possible influence of tree size on PD and FD with a linear model respectively, which included the log+1 transformed DBH and forest type as predictors. To remove the shared information of FD and PD, we decoupled the signal of traits from the phylogeny (decoupled FD) by using the approach described in de Bello et al. (2017). Further, for a detailed picture of the functional changes associated with human disturbance, we calculated CWMs of individual traits on tree scale using the package ‘FD’ (Laliberté et al., 2010). On tree scale, we compared then the SES of FD and PD, and the CWMs, between primary and secondary forests with Kruskal-Wallis tests (package ‘ggpubr’; Kassambara, 2020).

Finally, we classified non-native species based on the Global Ant Biodiversity Informatics (GABI) database (Guénard et al., 2017). To test their effect on the SES of PD and FD, and on CWMs measures, we excluded them from our occurrence data and applied the same analysis procedure as above.

## Results

### Community structure and taxonomic and phylogenetic diversity

We sampled 852 trees, with a total of 128 ant species. Both forests were fundamentally different in their vegetation structure and ant assemblages (Figs. 1 and 2). The arboreal ant communities were phylogenetically diverse, representing 39 genera from 7 subfamilies (Fig. 2). The secondary forest lacked dorylines and ectatommines, and ponerines were far rarer than in the primary forest (only a single occurrence; Fig. 2). 31 (24%) species occurred in both forests but had different abundances (Fig. 2). Non-native ant species were found in both forests (10 species in total) but were much more common in secondary forest than in primary forest (28% and 1% of species occurrences respectively; Fig. 2; Table 1). The species overlap between nesters and foragers was higher in the primary forest (Bray-Curtis dissimilarity 0.55) than in the secondary (Bray-Curtis dissimilarity 0.70).

**Figure 1.**
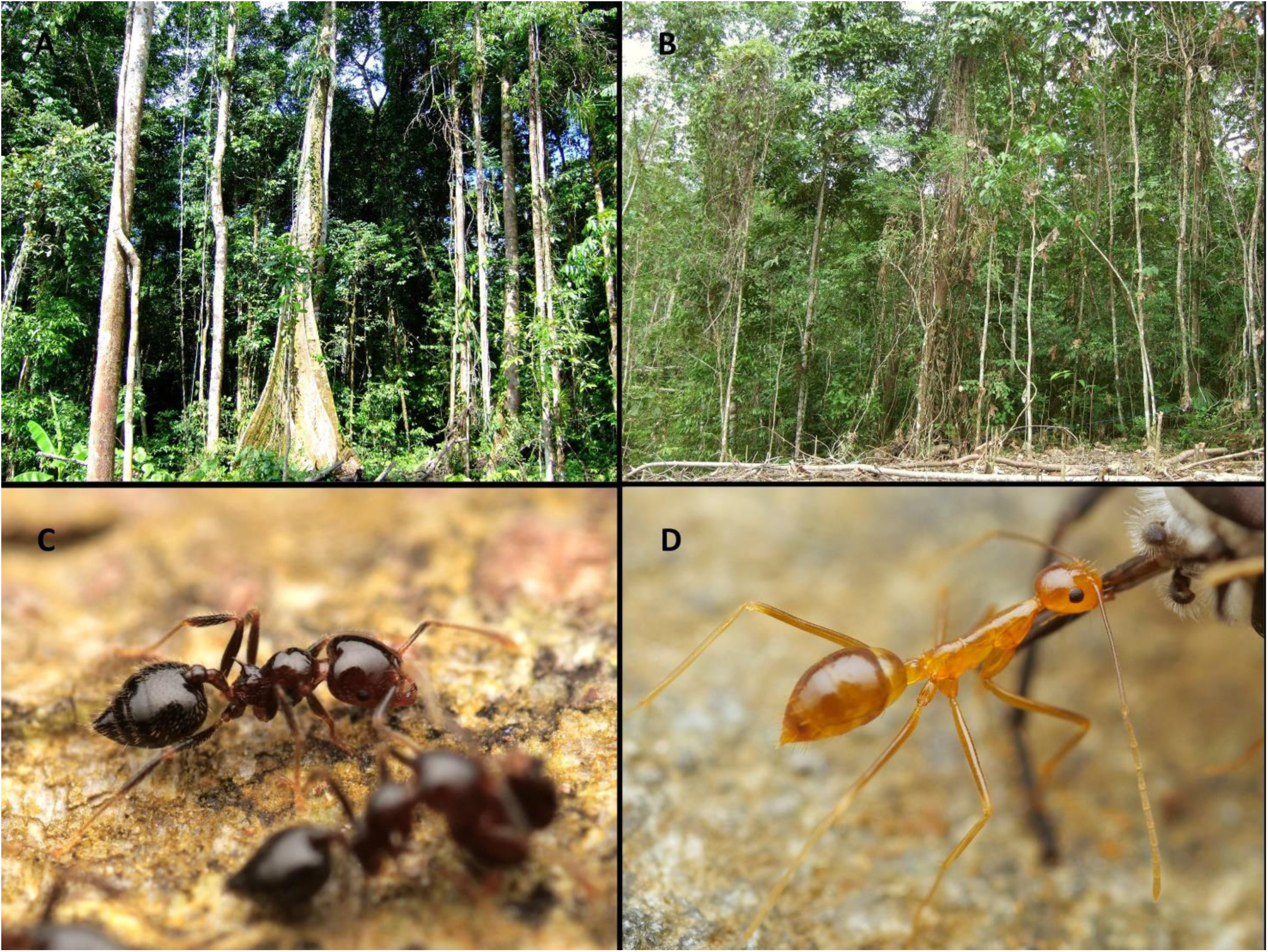
Images of the sampled primary (A) and secondary forest (B). The primary forest has larger trees and is overall more complex with more epiphytic growth. The most common ant in primary forest trees is the acrobat ant *Crematogaster polita* (A), a supercolonial ant species that builds carton nests in the canopy of trees. Conversely, the most common ant of the secondary forest is the globally invasive Yellow Crazy Ant *Anoplolepis gracilipes* (D), which has large colonies that nest on the ground and only forage into the tree canopy. Note the strong differences of functional traits just between these two most species: *Anoplolepis gracilipes* has longer legs and antenna, as well as much larger body and eye size, and no spines. Image credit: A,B: Petr Klimes, C,D: Philipp Hoenle.

**Figure 2.**
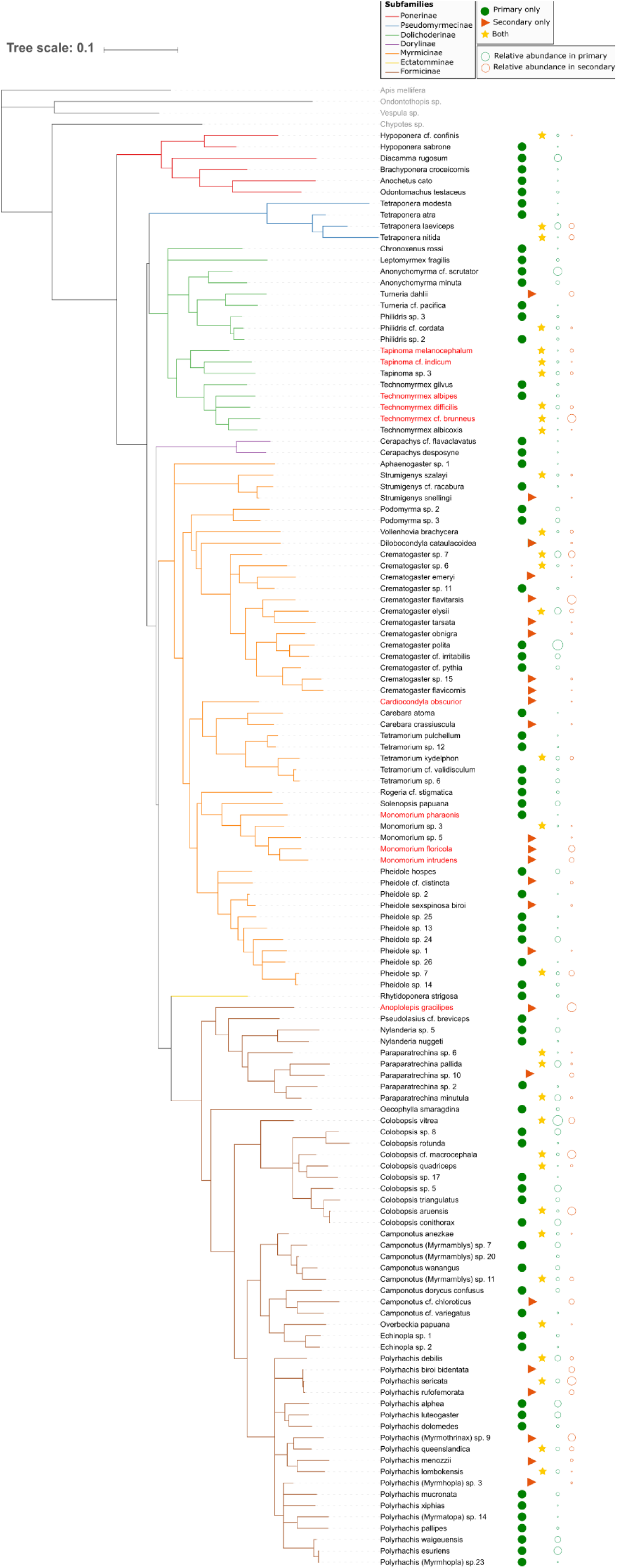
Bayesian phylogeny of arboreal ant communities sampled in 0.4 ha of primary and 0.4 ha of secondary lowland rainforest in Papua New Guinea (127 ant species across both forest types), and their relative abundance in each forest community. Outgroup species are denoted in grey, and non-native species are in red. Clades are colour coded by subfamily. Symbols indicate in which forest type each of the species occurred: circles = primary forest only (n=70), triangles = secondary forest only (n=26), stars = present in both forests (n=31). Open circles scaled by size indicate the relative abundance of the species in each forest (i.e., number of occupied trees), on a log scale.

In both plots over 90% of trees contained ants, with a mean of 3.3 species per tree, respectively. Primary forest had almost double the ant species richness at the plot level compared to secondary forest (Table 1). On tree scale, richness was also higher in primary than secondary forest in all species and visitors, but was similar among nesters (Fig. 3F, Table 1).

**Figure 3.**
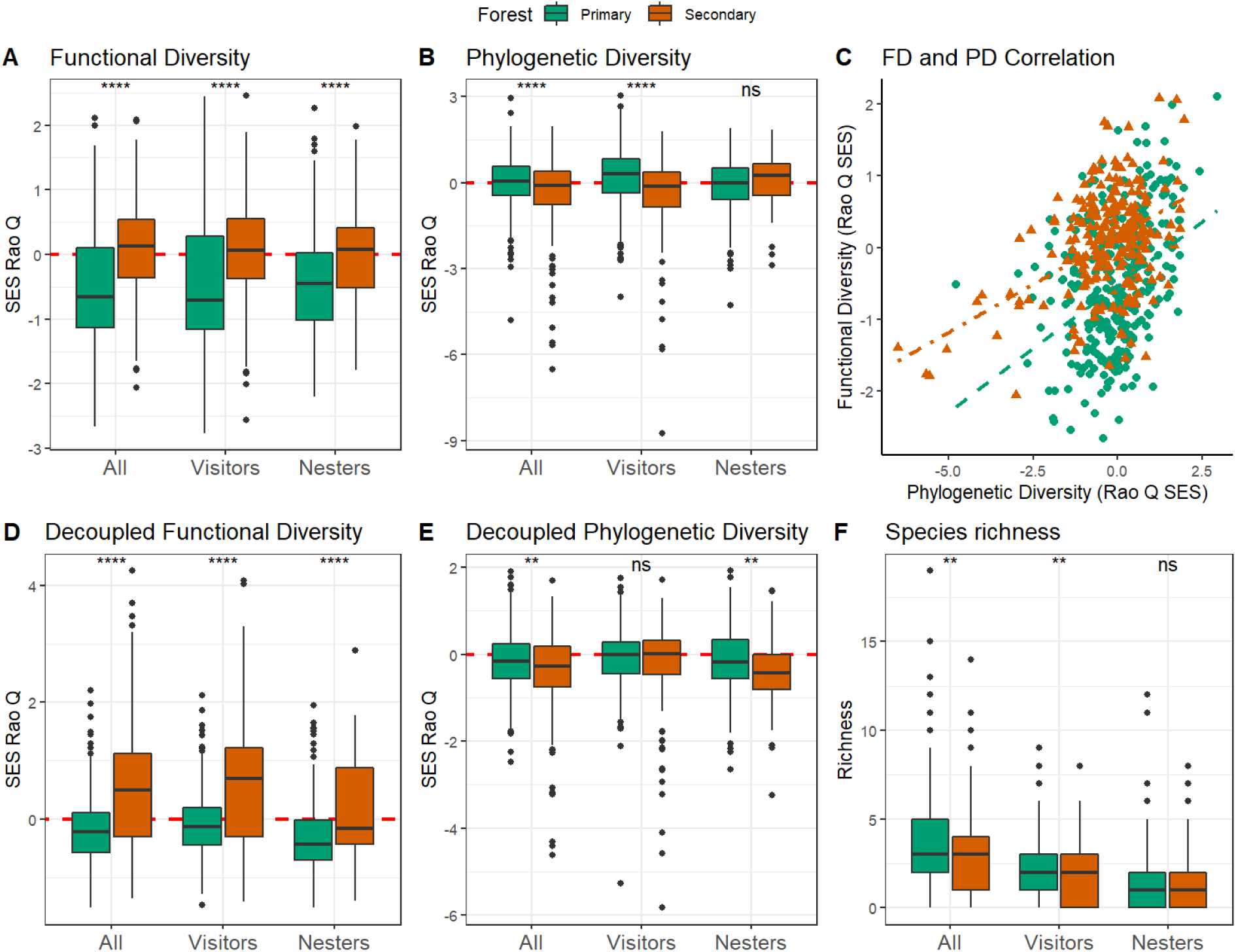
Arboreal ant functional diversity (A), phylogenetic diversity (B), the correlation between functional and phylogenetic diversity (C), the decoupled functional (D) and phylogenetic diversity (E) and the species richness (F) on tree scale. Box-plots show median values per a tree with 25–75% quartiles and with whiskers representing 1.5 interquartile ranges for all species combined (All), for foraging species not nesting in a focal tree (Visitors) and nesting species (Nesters). Both forest types are shown in different colours (primary forest – green; secondary forest – orange) or symbols (in C: triangles denote primary forest, circles secondary forest). The values for individual ant communities are compared between the two forest types with a Kruskall-Wallis test, where stars indicate statistically significant differences (*** p < 0.001, ** p < 0.01, * p 0.05, ns p > 0.05). Note that standardised effect sizes (i.e., Rao Q SES) are compared in A, B, D, and E where the dash horizontal line at 0 indicates random communities, while positive SES values indicate community overdispersion and negative values clustering. For mean observed and SES values and for tests of statistical significance of the SES against null distribution, see Table 1.

On the plot scale, secondary forest communities had lower PD, and were phylogenetically less dispersed than primary forests (except in nesters), but did not differ significantly from the null model expectation (Table 1). On tree scale, the PD of ants in the secondary forest was significantly clustered and in the primary forest significantly overdispersed (except in nesters; Fig. 3B, Table 1). These patterns were predominantly caused by the visitors, and not by nesters. These PD results were robust to the removal of non-native ants (Supplement Figure S1, Table S1).

### Functional diversity and trait differences

In contrast to PD patterns, FD was higher in secondary than in primary forests (Fig. 3A). Notably, the increased FD in secondary compared to primary forest was consistent on both plot and tree scales (Table 1). On plot scale, the primary forest was more functionally clustered than the secondary forest, but neither forest showed significant clustering or overdispersion in comparison to the null model (Table 1A). On tree scale, FD in primary forest was significantly clustered and in secondary forest overdispersed except for nesters and visitors (Table 1B; Fig. 3A). The pattern of higher FD in the secondary forest were robust for community subsets (Table 1B; Fig. 3A), and remained if non-native species were removed from the data set (Supplement Figure S2A, Table S1). Although FD and PD values were highly correlated with each other in both forest types (spearman rank correlation, p-value <0.001, ρ = 0.263), we found that intercept values were significantly higher in secondary forest (linear model intercept = 0.160±0.041) than in primary (linear model intercept= −0.522±0.041), with no significant interaction with forest type (linear model, p= 0.105, Fig. 3C).

Phylogenetic signal was present in all traits (Table S2). After decoupling the phylogeny from the traits, null model comparisons continued to indicate random dispersion for all forest types and communities (Table 1A).

However, on tree level, the secondary forest FD showed a stronger overdispersion and increase in FD, compared to primary forest that was significantly clustered (Fig. 3D, Table 1B). The PD, on the other hand, was barely affected by decoupling and lead to overall smaller differences between primary and secondary forests (Fig. 2E). The decoupled values overall aligned with previously patterns and showed for the most part significant deviation from zero distribution in almost all communities (except for the FD of secondary nesters and the PD of visitors Fig. 3D, Table 1B). Decoupled diversity results were for the most part stable to the exclusion of non-native ants in the whole community (Supplement Fig. S2, Table S1). Neither FD nor PD was significantly impacted by DBH (linear models, FD: R2=0.10, F=37.79, df= 659, p=0.28; PD: R2=0.04, F=14.61, df= 659, p=0.95).

With the exception of head length (a proxy of body size), community-weighted means differed between primary and secondary forests in whole tree communities (community matrix ‘all’). Primary forests ant communities were more polymorphic, had longer mandibles, higher eye positioning, larger clypeus, and broader heads, and more cuticular sculpturing. Secondary forest ant communities had higher spinosity, larger eyes and longer legs (Fig. 4). Several traits differed between the two forests also in nesters and visitors, leading to a different contribution of each functional group to some of the above whole-communities patterns (e.g., higher spinosity and longer legs in secondary forest were driven by visitors, Fig. 4). Notably, head lengths, a surrogate of body size, of visitors were smaller but of nesters larger in primary than secondary forest, while in the whole community this influence was masked, as there was no difference in body size in ‘all’ ants (Fig. 4A). Overall, the difference between nesters and visitors tended to be larger within secondary than within primary forest in most traits (Fig. 4).

**Figure 4.**
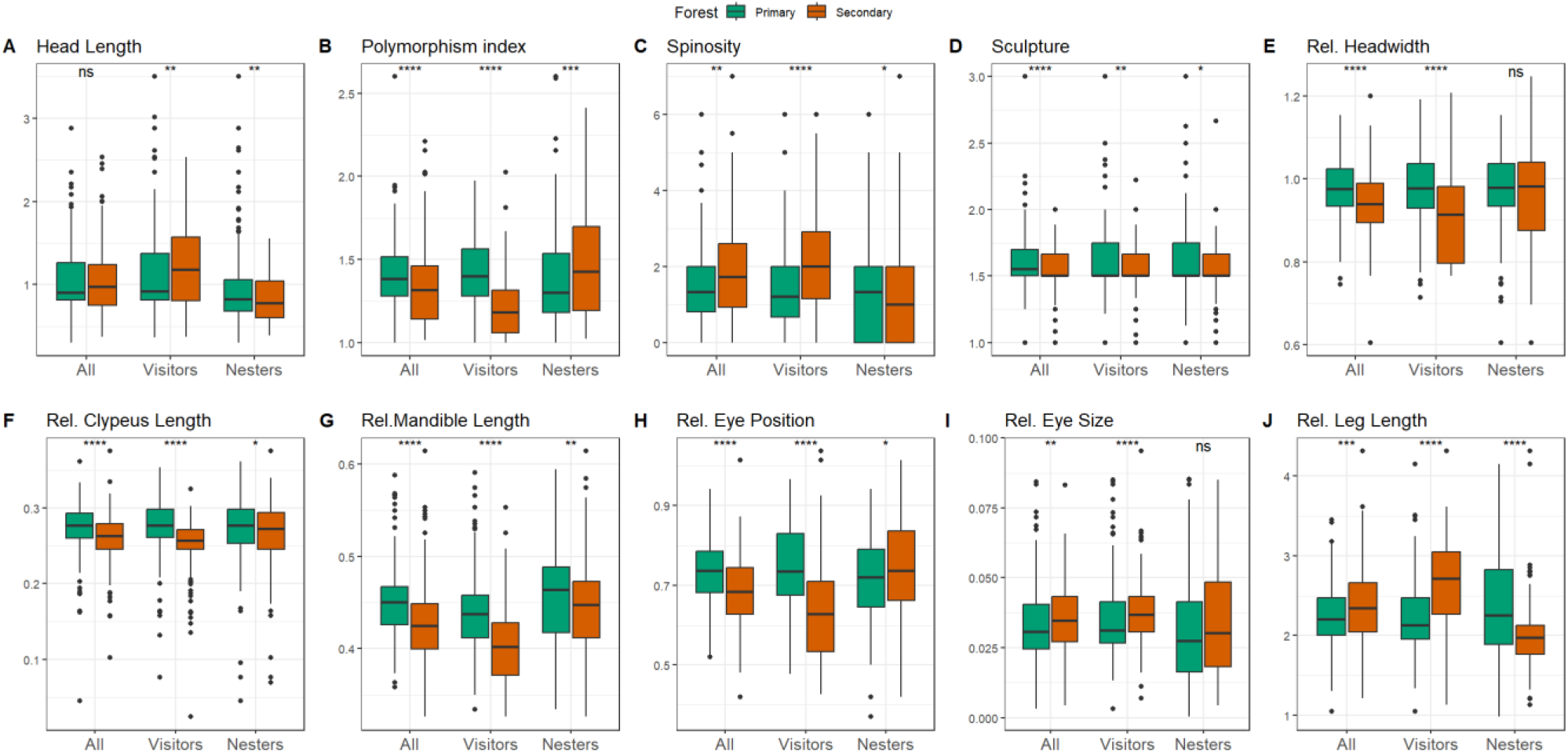
Community weighted means of all used ant traits (tree scale). Box-plots show median values per a tree with 25–75% quartiles and with whiskers representing 1.5 interquartile ranges for all ant species occurrences (All), for ant occurrences that were foraging but not nesting in a focal tree (Visitors) and for only nesting ant occurrences for each tree (Nesters). Both forest types are shown in different colours (primary forest – green; secondary forest – orange) and the values for individual ant communities are compared between the two forest types with a Kruskall-Wallis test. Stars indicate statistically significant differences (*** p < 0.001, ** p < 0.01, * p 0.05, ns – p > 0.05).

## Discussion

*Disturbance effects on functional and phylogenetic diversity of the ant communities* Disturbance results in profound changes in biological communities, usually diminishing species diversity accompanied with a simultaneous decrease in ecosystem functioning and functional diversity (Naeem et al., 2015: Ewers et al., 2015, but see Sreekar et al., 2021). Here, we show that the functional structure of an ecologically important group, arboreal ants, does not adhere to this pattern. In contrast to our initial expectation, FD weakly increased after disturbance since about 10 years of forest succession on a previously slash-and-burn garden. Still, the primary forest hosted a greater number of ant species, which were more diverse in their evolutionary history (PD). Our finding contrasts with several studies which documented a decrease in both FD and PD under human disturbance, including ants (Bihn et al., 2010; Letcher et al., 2012; Whitfeld et al., 2012; Mo et al., 2013; Bu et al., 2014).

The decrease in taxonomic diversity in secondary forests was previously well explained by changes in forest vegetation structure (Klimes et al., 2012). We therefore expected that the less variable microclimate and less complex secondary vegetation (e.g., lack of epiphytes and hollow live branches, see Klimes 2017) would select ant communities of similar (i.e., clustered) traits. We found this hypothesis not to be supported. Instead, we hypothesize that competitive interactions between species in secondary forest leads to the co-occurrence of morphologically more dissimilar species (i.e., overdispersion). The trait overdispersion was particularly strong among related species, as can be gasped by the decoupled functional diversity patterns: Even after correcting for phylogeny, the primary forest communities were morphologically more clustered than the secondary forest, hinting at a stronger role of environmental filtering for community assembly. Furthermore, our results suggest that most of that increased trait divergence among closely related secondary forest species occurs at the level of co-foraging species from surrounding trees (visitors), but much less so in the nesters. Only few previous faunistic studies showed functional overdispersion in more disturbed habitats, for instance in recovering bird communities (Sreekar et al., 2021) and in island ants (Agavekar et al., 2018).

Plot scale analysis indicated random functional as well as phylogenetic structure in both forests, i.e. there were no significant deviation from the null models. In contrast, tree scale found in almost all cases significant deviations from 0 (i.e., indicating clustering or overdispersion). This is not surprising because the taxa-swap null model is a much more conservative test (Hardy, 2008; Götzenberger et al., 2016). To reconcile these results, we conclude that secondary forest ant communities are more overdispersed in FD and clustered in PD in comparison to primary forest communities on both scales, but they do not necessarily reflect a strongly clustered or dispersed community *per-se*.

The interpretation of functional and phylogenetic dispersion patterns is not straight-forward, since mechanisms other than environmental filtering can lead to clustering (e.g. colonisation; Li et al., 2015). In the case of arboreal ants there is strong prior evidence that competition plays an important role in community assembly, as highly competitive ant species are known to control large territories and tend to exclude each other (so called ant mosaics; Mottl et al. 2021). Therefore, under the assumption that competition is increased through disturbance (Fayle et al., 2013), our observations are consistent with the hypothesis that competition leads to trait overdispersion in secondary forests. Our results fits well to the Interaction Hypothesis outlined by Prinzing et al. (2008), which predicts that competition is stronger among closely related species since traits are more likely to be used for similar resources, and thus a phylogenetically more related assemblage can lead to (either through evolutionary time or ecological interactions) higher functional diversity.

### Disturbance effects on traits of the ant communities

In our system, the impacts of forest disturbance on ant morphology were ubiquitous: When we considered the mean functional traits, all differed between primary and secondary forest, with the exception of body size. Thermal tolerance, competitive interactions, and diet likely play important roles in structuring ant communities in arboreal habitats (Blüthgen et al., 2003; Kaspari et al., 2015). Our findings largely support these assembly mechanisms. The changes are most evident in functional traits associated with diet: Traits associated with predatory behaviour were more prevalent in primary forest (e.g. relatively broader heads, longer mandibles, higher eye positioning, smaller eye size; Gibb et al., 2015; Parr et al., 2017). However, the clypeus, a structure involved in sucking ability and indicative of a low trophic level diet of sugar resources (Davidson et al., 2004), was also longer in the primary forest. Intraspecific polymorphism, which is associated with larger colony sizes and division of labour, was higher in primary forest: This could be an indication of higher competition in primary forest, which contrasts with the finding of more functional clustering among traits. Similarly, cuticular sculpturing was more common in primary forests, making species more robust to microclimatic changes or predation (Buxton et al., 2021). Finally, the spinosity, a defensive trait, is higher in secondary forests, potentially indicating higher levels of vertebrate predation and/or higher competition between the ants (Blanchard et al., 2020).

Our study is the first that examined the functional contribution of visiting and nesting species in tree canopies. As expected, there was less species overlap among visitors and nesters in secondary than in primary forest. One likely explanation is that in secondary forests, the trees are not as large and do not provide as many nesting resources as in primary forests. Trees are thus more shared between arboreal nesters and foragers that either nest mostly on the ground (e.g., the invasive *Anoplolepis gracilipes*), or forage from other trees. However, the functional diversity of secondary forests is higher even when visitors were disregarded, hence the contribution of visiting species to overall FD and PD was negligible. This could be explained by a low contribution of the species exclusively nesting on the ground, as the species not building any nests in our plots were rare, and both ant communities were dominated by arboreal fauna (see Klimes et al., 2015).

However, when considering individual traits, the differences between visitors and nesters became more evident. For instance, the longer average leg length in secondary forests is mainly driven by visitors - in fact, if only nesters are considered, primary forests communities have longer legs. One reason for the increase is the abundant invasive yellow crazy ant *Anoplolepis gracilipes*, a ground-nesting ant which builds only occasionally satellite worker nests on trees and has comparatively long legs. Other community level trait means where the differences between forest types emerge because of visitors are spinosity, polymorphisms, the head width, and eye positioning. While these differences are not captured in the functional diversity index, they suggest a more diverse or at least different functional contribution of nesters and visitors after disturbance.

### Effects of non-native ant species

Non-native species can replace native species and perform key ecosystem functions worse (Goodenough, 2010; Gallardo et al., 2016; Wong et al., 2020; Wong et al. 2021). Disturbance facilitates the spread of non-native species, and in our case invasive species reached high abundances in the secondary forest (28% of total species occurrences). However, despite this strong impact by numbers alone, neither FD nor PD was sensitive to the exclusion of non-native ants from the analysis. Only the species richness decreased on tree scale, and phylogenetic clustering in secondary forest was increased. Such changes are expected, however, if a large proportion of the community is excluded. The robustness of the PD and TD to non-native ant exclusion suggests a surprising resistance of the local ecosystem to the non-native species, contrasting other findings with invasive fire ants which reduced FD (Wong et al., 2020; Wong et al. 2021). However, manipulative experiments or time series are needed to clearly disentangle the effects of non-native species from habitat disturbance.

### Correlation of phylogenetic and functional diversity

Despite having a lower overall FD but higher PD in primary forests, community level PD and FD were still positively correlated in both forest types. The pattern is consistent with most correlations reported in the literature (see Cadotte et al. 2019), since many ecologically-relevant traits display a phylogenetic signal (Gerhold et al., 2015). In a review of PD and FD correlation, Cadotte et al. (2019) reported that roughly half of the ecological studies found incongruent patterns of FD and PD.

In our case study, the secondary forest hosts functionally richer but phylogenetically poorer ant communities. It is unclear if either the FD or the PD provide a more accurate picture of the arboreal ants’ contribution to ecosystem functioning. Although the investigated morphological traits relate to various ecological roles (Parr et al., 2017), we are still far away from a comprehensive functional perspective. Hence, we cannot exclude the possibility that other physiological and behavioural traits that we did not measure (such as colony size or intraspecific aggression) may be more important in these communities and distorted our results. Finally, PD may convey only weak ecological signals, since the ants of New Guinea likely are the result of an evolutionary recent and fast population differentiation and species radiation (5– 10 mya; e.g., Janda et al., 2016), similar to other insects (Toussaint et al., 2014). In fact, there is only one endemic ant genus known from the island (*Anciridris* – not present in our study), while both forests we studied had abundant lineages which showed recent diversification (e.g., species of *Camponotus* and *Polyrhachis*). Thus, closely related species of ants might show stronger morphological and ecological separation than is typical for other areas, lowering the sensitivity of our PD index.

## Conclusion

We quantified the response of functional and phylogenetic diversity in a tropical rainforest of rarely studied canopy ants, an insect guild of a high ecological importance and biomass (Davidson et al., 2003). We conclude that disturbance in this forest lead to increased FD, but to a phylogenetically and taxonomically less diverse arboreal ant fauna. The overdispersed traits in secondary forest are likely the result of higher competition in this less structurally diverse vegetation. Notably, the increased FD and a stronger trait overdispersion after decoupling it from PD signal is evidence that this unexpected increase occurs in closely related species. The New Guinea lowlands contains one of the most diverse rainforests in the world (Novotny et al., 2010), and although our study sampled complete arboreal ant communities from relatively large forest plots, it compares just two forest plots, and a larger sampling effort across more areas is needed to generalize our findings.

Interestingly, the introduction of non-native species and disturbance did not result in a reduction of the ants’ functional diversity in the canopy. While we did not assess any measure of ecosystem functioning directly, this will be of paramount importance for future investigation. Our study underlines the need of gathering both functional and phylogenetic data – had we investigated only one of them at a time, we would have come to opposite conclusions regarding their ecological implications. Our result urge for caution in the interpretation of trait diversity after forest disturbance, as a functionally more diverse community might not necessarily reflect better ecosystem health.

## Acknowledgements

We thank Filip Damen and the local community of Wanang who allowed us to work in their forest and logistically supported the project, and to Prof. Vojtech Novotny and Prof. George Weiblen for designing and supporting the felling project. We are grateful to the staff of New Guinea Binatang Research Center for their assistance in the field with sample collection and logistics. We thank Rudy Kohout, Archie MacArthur, Milan Janda, Eli Sarnat, Steven O. Shattuck and Phil Ward for helping to identify the ant species. We are grateful to Kate Parr for advice on functional measures. We thank Michaela Borovanska for advice on molecular protocols, and Ondrej Mottl, Alena Bartonova, and Jan Zima for help with DNA isolations and preparation of the specimens for sequencing. This work was supported by Czech Science Foundation Standard project (21-00828S), European Research Council (Project no. 669609) and the Grant Agency of University of South Bohemia (GAJU; 152/2016/P, 038/2019/P).

## Author contributions

POH, NSP and PK conceived the study. PK, CI and MR led the field work and collected the data. PK and ML sorted and identified the ant samples. NSP and ML measured the functional traits. NSP and PMM assembled the DNA sequences. PMM aligned the sequences and constructed the ant phylogeny and NSP finalised the phylogeny figure. PK and POH assembled the data for analyses. POH and PK conceived the analyses with further input from FB and TB. POH wrote the code and analysed the data. POH, PK and NSP wrote the manuscript. All authors have contributed critically to the drafts and gave final approval for publication of the submitted manuscript.

## Data availablity statement

All data and scripts underlying this manuscript will be publicly available in an appropriate repository (e.g., dryad).

## References

Agavekar, G., Garcia, F. H., & Economo, E. P. (2017) Taxonomic overview of the hyperdiverse ant genus Tetramorium Mayr (Hymenoptera, Formicidae) in India with descriptions and X-ray microtomography of two new species from the Andaman Islands. PeerJ, 5, e3800.

Andersen, A.N. (2000) The ants of northern Australia: A guide to the monsoonal fauna. CSIRO.

Basset, Y., Cizek, L., Cuénoud, P., Didham, R. K., Novotny, V., Ødegaard, F., … & Leponce, M. (2015) Arthropod distribution in a tropical rainforest: tackling a four dimensional puzzle. PLOS ONE, 10(12), e0144110.

Parr, C. L., & Bishop, T. R. (2022) The response of ants to climate change. Global Change Biology, 28, 3188–3205.

Bihn, J.H., Gebauer, G., & Brandl, R. (2010) Loss of functional diversity of ant assemblages in secondary tropical forests. Ecology, 91, 782–92.

Blanchard, B. D., Nakamura, A., Cao, M., Chen, S. T., & Moreau, C. S. (2020) Spine and dine: A key defensive trait promotes ecological success in spiny ants. Ecology and evolution, 10(12), 5852–5863.

Blüthgen, N., Gebauer, G., & Fiedler, K. (2003) Disentangling a rainforest food web using stable isotopes: dietary diversity in a species-rich ant community. 426–435.

Blomberg, S. P., Garland Jr, T., & Ives, A. R. (2003) Testing for phylogenetic signal in comparative data: behavioral traits are more labile. Evolution, 57(4), 717–745.

Bolton, B. (1994) Identification Guide to the Ant Genera of the World. Harvard University Press,

Bu, W., Zang, R., & Ding, Y. (2014) Functional diversity increases with species diversity along successional gradient in a secondary tropical lowland rainforest. Tropical Ecology, 55, 393–401.

Buxton, J. T., Robert, K. A., Marshall, A. T., Dutka, T. L., & Gibb, H. (2021) A cross-species test of the function of cuticular traits in ants (Hymenoptera: Formicidae). Myrmecological News, 31, 31–46.

Cadotte, M. W., Carboni, M., Si, X., & Tatsumi, S. (2019) Do traits and phylogeny support congruent community diversity patterns and assembly inferences? Journal of Ecology, 107(5), 2065–2077.

Cadotte, M.W., Jonathan Davies, T., Regetz, J., Kembel, S.W., Cleland, E., & Oakley, T.H. (2010) Phylogenetic diversity metrics for ecological communities: Integrating species richness, abundance and evolutionary history. Ecology Letters, 13, 96–105.

Cavender-Bares, J., Kozak, K.H., Fine, P.V.A., & Kembel, S.W. (2009) The merging of community ecology and phylogenetic biology. Ecology Letters, 12, 693–715.

Chaves, L. F.. (2010). An entomologist guide to demystify pseudoreplication: data analysis of field studies with design constraints. Journal of medical entomology, 47(3), 291–298.

Davidson, D.W., Cook, S.C., & Snelling, R.R. (2004) Liquid-feeding performances of ants (Formicidae): ecological and evolutionary implications. Oecologia, 139, 335–335.

Davidson, D.W., Cook, S.C., Snelling, R.R., & Chua, T.H. (2003) Explaining the Abundance of Ants in Lowland Tropical Rainforest Canopies. 300, 969–972.

de Bello, F. (2012) The quest for trait convergence and divergence in community assembly: are null-models the magic wand? Global Ecology and Biogeography, 21(3), 312–317.

de Bello, F., Carmona, C. P., Lepš, J., Szava-Kovats, R., & Pärtel, M. (2016) Functional diversity through the mean trait dissimilarity: resolving shortcomings with existing paradigms and algorithms. Oecologia, 180(4), 933–940.

de Bello, F., Šmilauer, P., Diniz-Filho, J. A. F., Carmona, C. P., Lososová, Z., Herben, T., & Götzenberger, L. (2017) Decoupling phylogenetic and functional diversity to reveal hidden signals in community assembly. Methods in Ecology and Evolution, 8(10), 1200–1211.

de Bello, F., Carmona, C. P., Dias, A. T., Götzenberger, L., Moretti, M., & Berg, M. P. (2021) Handbook of trait-based ecology: From theory to R tools. Cambridge University Press.

Diaz, S. & Cabido, M. (2001) Vive la difference: plant functional diversity matters to ecosystem processes. Trends in Ecology & Evolution, 16, 646–655. https://doi.org/10.1016/s0169-5347(01)02283-2

Dunn, R.R. (2004) Recovery of Faunal Communities During Tropical Forest Regeneration. Conservation Biology, 18, 302–309.

Ewers, R.M., Boyle, M.J.W., Gleave, R.A., et al. (2015) Logging cuts the functional importance of invertebrates in tropical rainforest. Nature Communications, 6, 1–7.

Fayle, T. M., Turner, E. C., & Foster, W. A. (2013) Ant mosaics occur in SE Asian oil palm plantation but not rain forest and are influenced by the presence of nest-sites and non-native species. Ecography, 36(9), 1051–1057.

Floren, A., Wetzel, W., & Staab, M. (2014) The contribution of canopy species to overall ant diversity (Hymenoptera: Formicidae) in temperate and tropical ecosystems. Myrmecological News, 19(1), 65–74.

Flynn, D. F., Mirotchnick, N., Jain, M., Palmer, M. I., & Naeem, S. (2011) Functional and phylogenetic diversity as predictors of biodiversity–ecosystem-function relationships. Ecology, 92(8), 1573–1581.

Gallardo, B., Clavero, M., Sánchez, M. I., & Vilà, M. (2016) Global ecological impacts of invasive species in aquatic ecosystems. Global change biology, 22(1), 151–163.

Gerhold, P., Cahill Jr, J. F., Winter, M., Bartish, I. V., & Prinzing, A. (2015) Phylogenetic patterns are not proxies of community assembly mechanisms (they are far better). Functional Ecology, 29(5), 600–614.

Gibb, H., Dunn, R.R., Sanders, N.J., et al. (2017a) A global database of ant species abundances. Ecology, 98, 883–884.

Gibb, H., Sanders, N.J., Dunn, R.R., Arnan, X., Vasconcelos, H.L., Donoso, D.A., Andersen, A.N., Silva, R.R., Bishop, T.R., Gomez, C., Grossman, B.F., Yusah, K.M., Luke, S.H., Pacheco, R., Pearce-Duvet, J., Retana, J., Tista, M., & Parr, C.L. (2017b) Habitat disturbance selects against both small and large species across varying climates. Ecography, 1–9.

Gibb, H., Stoklosa, J., Warton, D.I., Brown, A.M., Andrew, N.R., & Cunningham, S.A. (2015) Does morphology predict trophic position and habitat use of ant species and assemblages? Oecologia, 177, 519–531.

Götzenberger, L., Botta-Dukát, Z., Lepš, J., Pärtel, M., Zobel, M., & de Bello, F. (2016) Which randomizations detect convergence and divergence in trait-based community assembly? A test of commonly used null models. Journal of Vegetation Science, 27(6), 1275–1287.

Goodenough, A. E. (2010) Are the ecological impacts of alien species misrepresented? A review of the “native good, alien bad” philosophy. Community Ecology, 11, 13–21.

Guénard, B., Weiser, M., Gomez, K., Narula, N., & Economo, E.P. (2017) The Global Ant Biodiversity Informatics (GABI) database: a synthesis of ant species geographic distributions. Myrmecological News, 24, 83–89.

Guilherme, D. R., Souza, J. L. P., Franklin, E., Pequeno, P. A. C. L., das Chagas, A. C., & Baccaro, F. B. (2019) Can environmental complexity predict functional trait composition of ground-dwelling ant assemblages? A test across the Amazon Basin. Acta Oecologica, 99, 103434.

Hansen, M.C.C., Potapov, P. V, Moore, R., et al. (2013) High-resolution global maps of 21st-century forest cover change. Science, 342, 850–854.

Hardy, O. J. (2008) Testing the spatial phylogenetic structure of local communities: statistical performances of different null models and test statistics on a locally neutral community. Journal of ecology, 96(5), 914–926.

Hethcoat, M. G., B. J. King, F. F. Castiblanco, C. M. Ortiz-Sepúlveda, F. C. P. Achiardi, F. A. Edwards, C. Medina, J. J. Gilroy, T. Haugaasen, & Edwards, D. P. (2019) The impact of secondary forest regeneration on ground-dwelling ant communities in the Tropical Andes. Oecologia, 191, 475–482.

Hoenle, P. O., Donoso, D. A., Argoti, A., Staab, M., von Beeren, C., & Blüthgen, N. (2022) Rapid ant community reassembly in a Neotropical forest: Recovery dynamics and land-use legacy. Ecological Applications, e2559. https://doi.org/10.1002/eap.2559

Jucker, T., Hardwick, S. R., Both, S., Elias, D. M., Ewers, R. M., Milodowski, D. T., … & Coomes, D. A. (2018) Canopy structure and topography jointly constrain the microclimate of human-modified tropical landscapes. Global change biology, 24(11), 5243–5258.

Kaspari, M., Clay, N.A., Lucas, J., Yanoviak, S.P., & Kay, A. (2015) Thermal adaptation generates a diversity of thermal limits in a rainforest ant community. Global Change Biology, 21, 1092–1102.

Klimes P. (2017) Diversity and specificity of ant-plant interactions in canopy communities: insights from primary and secondary tropical forests in New Guinea. In: Ant-Plant Interactions: Impacts of Humans on Terrestrial Ecosystems, Cambridge University Press, p 26–51.

Klimes, P., Fibich, P., Idigel, C., & Rimandai, M. (2015) Disentangling the diversity of arboreal ant communities in tropical forest trees. PLOS ONE, 10, e0117853.

Klimes, P., Idigel, C., Rimandai, M., Fayle, T.M., Janda, M., Weiblen, G.D., & Novotny, V. (2012) Why are there more arboreal ant species in primary than in secondary tropical forests? Journal of Animal Ecology, 81, 1103–12.

Kraft, N.J.B., Adler, P.B., Godoy, O., James, E.C., Fuller, S., & Levine, J.M. (2015) Community assembly, coexistence and the environmental filtering metaphor. Functional Ecology, 29, 592–599.

Lach, L., Parr, C.L., & Abbott, K.L. (2010) Ant Ecology. Oxford University Press.

Lawton, J.H., Bignell, D.E., Bolton, B., Bloemers, G.F., Eggleton, P., Hammond, P.M., Hodda, M., Holt, R.D., Larsen, T.B., Mawdsley, N.A., Stork, N.E., Srivastava, D.S., & Watt, A.D. (1998) Biodiversity inventories, indicator taxa and effects of habitat modification in tropical forest. Nature, 391, 72–76.

Leponce, M., Dejean, A., Mottl, O., & Klimes, P. (2021). Rapid assessment of the three-dimensional distribution of dominant arboreal ants in tropical forests. Insect Conservation and Diversity, 14(4), 426–438.

Letcher, S.G., Chazdon, R.L., Andrade, A.C.S., Bongers, F., van Breugel, M., Finegan, B., Laurance, S.G., Mesquita, R.C.G., Martínez-Ramos, M., & Williamson, G.B. (2012) Phylogenetic community structure during succession: Evidence from three Neotropical forest sites. *Perspectives in Plant Ecology*, Evolution and Systematics, 14, 79–87.

Li, S. P., Cadotte, M. W., Meiners, S. J., Hua, Z. S., Jiang, L., & Shu, W. S. (2015) Species colonisation, not competitive exclusion, drives community overdispersion over long-term succession. Ecology Letters, 18(9), 964–973.

Liu, C., Guénard, B., Blanchard, B., Peng, Y.Q., & Economo, E.P. (2016) Reorganization of taxonomic, functional, and phylogenetic ant biodiversity after conversion to rubber plantation. Ecological Monographs, 86, 215–227.

Loiola, P. P., de Bello, F., Chytrý, M., Götzenberger, L., Carmona, C. P., Pyšek, P., & Lososová, Z. (2018) Invaders among locals: Alien species decrease phylogenetic and functional diversity while increasing dissimilarity among native community members. Journal of Ecology, 106(6), 2230–2241.

MacArthur, R., & Levins, R. (1967).The limiting similarity, convergence, and divergence of coexisting species. The american naturalist, 101(921), 377–385.

Martello, F., De Bello, F., Morini, M. S. D. C., Silva, R. R., Souza-Campana, D. R. D., Ribeiro, M. C., & Carmona, C. P. (2018) Homogenization and impoverishment of taxonomic and functional diversity of ants in Eucalyptus plantations. Scientific reports, 8(1), 1–11.

McAlpine, J.R., Keig, G., & Falls, R. (1983) Climate of Papua New Guinea. Commonwealth Scientific and Industrial Organization in association with the Australian National University Press, Canberra,

Mo, X.X., Shi, L.L., Zhang, Y.J., Zhu, H., & Slik, J.W.F. (2013) Change in Phylogenetic Community Structure during Succession of Traditionally Managed Tropical Rainforest in Southwest China. PLOS ONE, 8, 1–9.

Mottl, O., Plowman, N. S., Novotny, V., Gewa, B., Rimandai, M., & Klimes, P. (2019). Secondary succession has surprisingly low impact on arboreal ant communities in tropical montane rainforest. Ecosphere, 10(8), e02848.

Mottl, O., Yombai, J., Novotný, V., Leponce, M., Weiblen, G. D., & Klimeš, P. (2021). Inter-specific aggression generates ant mosaics in canopies of primary tropical rainforest. Oikos, 130(7), 1087–1099.

Naeem, S., Duffy, J.E., & Zavaleta, E. (2012) The Functions of Biological Diversity in an Age of Extinction. Science, 336, 1401–1406.

Novotny, V. (2010) Rain forest conservation in a tribal world: why forest dwellers prefer loggers to conservationists. Biotropica, 42(5), 546–549.

Novotny, V., Miller, S.E., Baje, L., Balagawi, S., Basset, Y., Cizek, L., Craft, K.J., Dem, F., Drew, R. a I., Hulcr, J., Leps, J., Lewis, O.T., Pokon, R., Stewart, A.J. a, Samuelson, G.A., & Weiblen, G.D. (2010) Guild-specific patterns of species richness and host specialization in plant-herbivore food webs from a tropical forest. The Journal of animal ecology, 79, 1193–203.

Oksanen, L. (2001) Logic of experiments in ecology: is pseudoreplication a pseudoissue?. Oikos, 94(1), 27–38.

Parr, C.L., Dunn, R.R., Sanders, N.J., et al. (2017) GlobalAnts: a new database on the geography of ant traits (Hymenoptera: Formicidae). Insect Conservation and Diversity, 10, 5–20.

Pfeiffer, M., Cheng Tuck, H., & Chong Lay, T. (2008) Exploring arboreal ant community composition and co-occurrence patterns in plantations of oil palm Elaeis guineensis in Borneo and Peninsular Malaysia. Ecography, 31(1), 21–32.

Prinzing, A., Reiffers, R., Braakhekke, W. G., Hennekens, S. M., Tackenberg, O., Ozinga, W. A., … & Van Groenendael, J. M. (2008) Less lineages–more trait variation: phylogenetically clustered plant communities are functionally more diverse. Ecology letters, 11(8), 809–819.

R Core Team (2014) R: A language and environment for statistical computing..

Rocha-Ortega, M., Arnan, X., Ribeiro-Neto, J.D., Leal, I.R., Favila, M.E., & Martínez-Ramos, M. (2018) Taxonomic and functional ant diversity along a secondary successional gradient in a tropical forest. Biotropica, 50, 290–301.

Sanders, N. J., Gotelli, N. J., Heller, N. E., & Gordon, D. M. (2003) Community disassembly by an invasive species. Proceedings of the National Academy of Sciences, 100(5), 2474– 2477.

Schmidt, C. A., & Shattuck, S. O. (2014) The higher classification of the ant subfamily Ponerinae (Hymenoptera: Formicidae), with a review of ponerine ecology and behavior. Zootaxa, 3817(1), 1–242.

Skarbek, C. J., Noack, M., Bruelheide, H., Härdtle, W., von Oheimb, G., Scholten, T., … & Staab, M. (2020) A tale of scale: Plot but not neighbourhood tree diversity increases leaf litter ant diversity. Journal of Animal Ecology, 89(2), 299–308.

Sreekar, R., Si, X., Sam, K., Liu, J., Dayananda, S., Goodale, U., … & Goodale, E. (2021) Land use and elevation interact to shape bird functional and phylogenetic diversity and structure: Implications for designing optimal agriculture landscapes. Journal of Applied Ecology, 58(8), 1738–1748.

Srivastava, D. S., Cadotte, M. W., MacDonald, A. A. M., Marushia, R. G., & Mirotchnick, N. (2012) Phylogenetic diversity and the functioning of ecosystems. Ecology letters, 15(7), 637–648.

Tschinkel, W. R. (2005) The nest architecture of the ant, Camponotus socius. Journal of Insect Science, 5(1), 9.

Toussaint, E. F., Hall, R., Monaghan, M. T., Sagata, K., Ibalim, S., Shaverdo, H. V., … & Balke, M. (2014) The towering orogeny of New Guinea as a trigger for arthropod megadiversity. Nature communications, 5(1), 1–10.

Tucker, C. M., Davies, T. J., Cadotte, M. W., & Pearse, W. D. (2018) On the relationship between phylogenetic diversity and trait diversity. Ecology, 99(6), 1473–1479.

Volf, M., Klimeš, P., Lamarre, G. P., Redmond, C. M., Seifert, C. L., Abe, T., … & Novotny, V. (2019) Quantitative assessment of plant-arthropod interactions in forest canopies: A plot-based approach. PLOS ONE, 14(10), e0222119.

Walker, J., & Stamps, J. (1986) A test of optimal caste ratio theory using the ant Camponotus (Colobopsis) impressus. Ecology, 67(4), 1052–1062.

Webb, C. O., Ackerly, D. D., McPeek, M. A., & Donoghue, M. J. (2002) Phylogenies and community ecology. Annual review of ecology and systematics, 33(1), 475–505.

Whitfeld, T.J.S., Kress, W.J., Erickson, D.L., & Weiblen, G.D. (2012) Change in community phylogenetic structure during tropical forest succession: evidence from New Guinea. Ecography, 35, 821–830.

Whitfeld, T. J.S., Lasky, J. R., Damas, K., Sosanika, G., Molem, K., & Montgomery, R. A. (2014) Species richness, forest structure, and functional diversity during succession in the New Guinea lowlands. Biotropica, 46(5), 538–548.

Whittaker, R.J., Willis, K.J., & Field, R. (2001) Scale and species richness: towards a general, hierarchical theory of species diversity. Journal of Biogeography, 28(4), 453–470.

Wong, M. K., Guénard, B., & Lewis, O. T. (2020) The cryptic impacts of invasion: functional homogenization of tropical ant communities by invasive fire ants. Oikos, 129(4), 585–597.

Wong, M. K., Tsang, T. P., Lewis, O. T., & Guénard, B. (2021) Trait-similarity and trait-hierarchy jointly determine fine-scale spatial associations of resident and invasive ant species. Ecography, 44(4), 589–601.

Xing, S., Leahy, L., Ashton, L. A., Kitching, R. L., Bonebrake, T. C., & Scheffers, B. R. (2023) Ecological patterns and processes in the vertical dimension of terrestrial ecosystems. Journal of Animal Ecology, 00, 1–14.

