## Supplementary Material for "Disturbance increases functional diversity but decreases phylogenetic diversity of an arboreal tropical ant community"

**Supplement Text 1.** Genomic DNA was extracted using the Genomic DNAKit Tissue (Geneaid Biotech Ltd., New Taipei City, Taiwan) following the manufacturer's protocol, and primers and laboratory protocols follow published studies (Folmer et al., 1994; Ward & Downie, 2005; Brady et al., 2006). Sequencing of both DNA strands was carried out by the company Macrogen (South Korea) and the edition and alignment of sequences were conducted in Geneious R6.1. Dataset alignment was carried out using the Clustal W algorithm (Thompson et al., 1994) with the following parameters: cost matrix with gap open cost 16 and gap extend cost 8. With the exception of one single individual of *Lordomyrma* sp. for which we could not obtain molecular data (and thus excluded it from all further analyses is this study), we assembled molecular data for all the 127 remaining species with workers. Specifically, we obtained COI and Wg sequences for 112 of 127 species, gathered missing COI and/or Wg for a further 4 species from the Genbank database, while the remaining 11 species were represented by a single gene (6 COI, 5 Wg; 8% of species). All ant specimens and DNA vouchers are deposited at the Institute of Entomology, Biology Centre of Czech Academy of Sciences, České Budějovice. All DNA sequences will be made available upon acceptance either in GenBank (Sample Numbers XY) or in BOLD (Barcode of Life) under the 'Ants of Papua New Guinea (ASPNA)' project.

The concatenated and single-gene datasets consisting of 127 taxa and partitioned by codon positions of COI and *wingless* were analysed in RAxML v8.1.11 (Stamatakis, 2014) and MrBayes v3.2.3 (Ronquist et al., 2012). Four Hymenopteran outgroup taxa were used to root the phylogeny: *Apis mellifera*, *Vespula* sp., *Chypotes* sp., and *Odontophotopsis* sp. We constrained the phylogenetic relationships among ant subfamilies following the comprehensive ant-family phylogeny of Moreau & Bell (2013). This approach was necessary because our molecular dataset consisted of two fast-evolving gene fragments, thus deep nodes were expected not to be fully resolved. Maximum Likelihood analyses in RAxML v8.1.11 were carried out under the 'rapid bootstrapping' algorithm with 1,000 iterations. We set the GTRGAMMA model for each partition and we searched for the best-scoring tree using the command '-f a'. We computed an extended majority rule consensus tree from 1,000 bootstraps trees using the command '-J'. Bayesian Inference through MrBayes v3.2.3 consisted of two independent runs each for 50 million generations, sampling every 5,000 generations. The nucleotide substitution scheme 'mixed + gamma' (Huelsenbeck et al., 2004) was set to each partition. We confirmed convergence of chains after applying a burn-in of 25% by checking that the final average standard deviation of split frequencies was below 0.01, PSRFs were close to 1.0, ESS values higher than 200, and log probabilities reached stationary distribution. We summarized the MCMC sampled trees using the 50% majority rule consensus approach. All phylogenetic analyses were carried out through CIPRES (Miller et al., 2010). To visualise the phylogeny with community data we uploaded data to iTOL v.4.1 and finally edited in Inkscape v.0.92. For further analyses, after checking the tree topology congruence between the RAxML and MrBayes outputs, we imported our concatenated MrBayes phylogeny into R v3.4.0, removed the outgroups and created a phylogenetic distance matrix using the function *cophenetic* implemented in the 'stats' package. The phylogeny was square root transformed before community analyses following the recommendation of Letten and Cornwell (2015).

### References


Brady, S. G., Schultz, T. R., Fisher, B. L., & Ward, P. S. (2006) Evaluating alternative hypotheses for the early evolution and diversification of ants. Proceedings of the National Academy of *Sciences*, **103**(48), 18172–18177.

Folmer, O., Black, M.B., Hoch, W., Lutz, R.A., & Vrijehock, R.C. (1994) DNA primers for amplification of mitochondrial cytochrome c oxidase subunit I from diverse metazoan invertebrates. *Molecular Marine Biology and Biotechnology*, **3**, 294–299.

Huelsenbeck, J. P., Larget, B., & Alfaro, M. E. (2004) Bayesian phylogenetic model selection using reversible jump Markov chain Monte Carlo. *Molecular biology and evolution*, **21**(6), 1123–1133.

Letten, A. D., & Cornwell, W. K. (2015). Trees, branches and (square) roots: why evolutionary relatedness is not linearly related to functional distance. Methods in Ecology and Evolution, 6(4), 439–444.

Miller, M. A., Pfeiffer, W., & Schwartz, T. (2010) Creating the CIPRES Science Gateway for inference of large phylogenetic trees. In: 2010 gateway computing environments workshop (GCE) (pp. 1–8). Ieee.

Moreau, C. S., & Bell, C. D. (2013) Testing the museum versus cradle tropical biological diversity hypothesis: phylogeny, diversification, and ancestral biogeographic range evolution of the ants. *Evolution*, **67**(8), 2240-2257.

Stamatakis, A. (2014) RAxML version 8: A tool for phylogenetic analysis and post-analysis of large phylogenies. *Bioinformatics*, **30**, 1312–1313.

Thompson, J. D., Higgins, D. G., & Gibson, T. J. (1994) CLUSTAL W: improving the sensitivity of progressive multiple sequence alignment through sequence weighting, position-specific gap penalties and weight matrix choice. *Nucleic acids research*, **22**(22), 4673–4680.

Ward, P. S., & Downie, D. A. (2005) The ant subfamily Pseudomyrmecinae (Hymenoptera: Formicidae): phylogeny and evolution of big-eyed arboreal ants. *Systematic Entomology*, **30**(2), 310–335.

**Table S1** Characteristics of arboreal ant communities sampled in 0.4 ha of primary and 0.4 ha of secondary lowland rainforest in Papua New Guinea, and of their taxonomic, functional (FD) and phylogenetic (PD) diversity, with non-native species excluded from the dataset. (A) values on plot scale and (B) values on tree scale. Standardised effect sizes from null models (Rao Q SES, see Methods) near zero indicate random communities, while positive SES indicate community overdispersion (highlighted as light blue) and negative SES indicate clustering (highlighted as light orange). On plot scale, we give the p-values in comparison to the taxa-swap null model, while for the tree scale we test the distribution of all SES values against 0 with a Wilcoxon's signed-rank test. Significant overdispersion and clustering values as well as their respective p-values are highlighted in bold (alpha = 0.05).

| **A) Plot scale** | Primary all | Secondary all | Primary visitors | Secondary visitors | Primary nesters | Secondary nesters |
|---|---|---|---|---|---|---|
| N of sampled trees | 472 | 380 | 472 | 380 | 472 | 380 |
| N of species occurrences | 1660 | 848 | 997 | 469 | 662 | 662 |
| Species richness | 95 | 49 | 60 | 29 | 80 | 42 |
| FD (Rao observed) | 0.192 | 0.205 | 0.188 | 0.201 | 0.209 | 0.200 |
| FD (Rao SES) | -1.211 | -0.284 | -0.923 | -0.044 | -1.294 | -0.428 |
| p value (null model) | 0.092 | 0.398 | 0.195 | 0.481 | 0.092 | 0.359 |
| decoupled FD SES | -0.303 | 0.975 | -0.008 | **1.904** | -1.039 | -0.743 |
| p value (null model) | 0.474 | 0.844 | 0.593 | **0.951** | 0.146 | 0.252 |
| PD (Rao Q observed) | 0.529 | 0.489 | 0.514 | 0.455 | 0.538 | 0.483 |
| PD (Rao Q SES) | -0.030 | -1.526 | 0.269 | -1.092 | -0.655 | -0.820 |
| p value (null model) | 0.529 | 0.053 | 0.659 | 0.120 | 0.250 | 0.196 |
| decoupled PD (Rao Q SES) | -0.338 | -1.566 | -0.222 | -1.486 | -0.686 | -1.128 |
| p value (null model) | 0.383 | 0.053 | 0.426 | 0.061 | 0.252 | 0.127 |

| **B) Tree scale** | Primary all | Secondary all | Primary visitors | Secondary visitors | Primary nesters | Secondary nesters |
|---|---|---|---|---|---|---|
| N of occupied trees | 442 | 314 | 362 | 229 | 324 | 239 |

| | | | | | | |
|---|---|---|---|---|---|---|
| N of trees with ≥ 2 species | 382 | 224 | 284 | 147 | 172 | 87 |
| mean species richness | 3.52 | 2.23 | 2.11 | 1.23 | 1.40 | 1.00 |
| FD (Rao Q obs.) | 0.142 | 0.147 | 0.132 | 0.135 | 0.130 | 0.137 |
| FD (Rao Q SES) | **-0.592** | **-0.198** | **-0.481** | -0.120 | **-0.418** | -0.059 |
| p value (against 0) | **<0.001** | **0.001** | **<0.001** | 0.458 | **<0.001** | 0.921 |
| mean decoupled FD (Rao Q SES) | **-0.162** | **0.572** | 0.033 | **1.019** | **-0.326** | **-0.327** |
| p values (against 0) | **<0.001** | **<0.001** | 0.322 | **<0.001** | **<0.001** | **<0.001** |
| mean PD (Rao Q obs.) | 0.408 | 0.347 | 0.372 | 0.307 | 0.344 | 0.325 |
| mean PD (Rao Q SES) | **0.088** | **-0.742** | **0.182** | **-0.665** | -0.144 | *-0.229* |
| p values (against 0) | **0.007** | **<0.001** | **<0.001** | **0.006** | 0.438 | *0.065* |
| mean decoupled SES PD | **-0.108** | **-0.839** | -0.063 | **-0.890** | **-0.136** | **-0.471** |
| p value (against 0) | **0.001** | **<0.001** | 0.271 | **<0.001** | **0.016** | **<0.001** |

**Table S2**. Blomberg's K of all traits used. Blomberg's K signal indicates how strong a trait is predicted by its phylogeny, with values ranging from 0 (no phylogenetic signal) to >2 (i.e., strongly phylogenetically conserved trait), under the assumption of the Brownian Motion model of evolution. In addition, we give p-values for the test if a trait has a significant phylogenetic signal, which is the case for all.

| Trait | Blomberg's K | p-value |
|---|---|---|
| Spinosity | 1.74 | **0.001** |
| Sculpture | 0.60 | **0.010** |
| Rel. Head width | 1.06 | **0.001** |
| Rel. Clypeus Length | 2.40 | **0.001** |
| Rel. Mandible Length | 1.15 | **0.001** |
| Rel. Eye Position | 1.19 | **0.001** |
| Rel. Leg Length | 1.03 | **0.001** |
| Head Length | 1.39 | **0.001** |
| Polymorphism index | 1.13 | **0.001** |

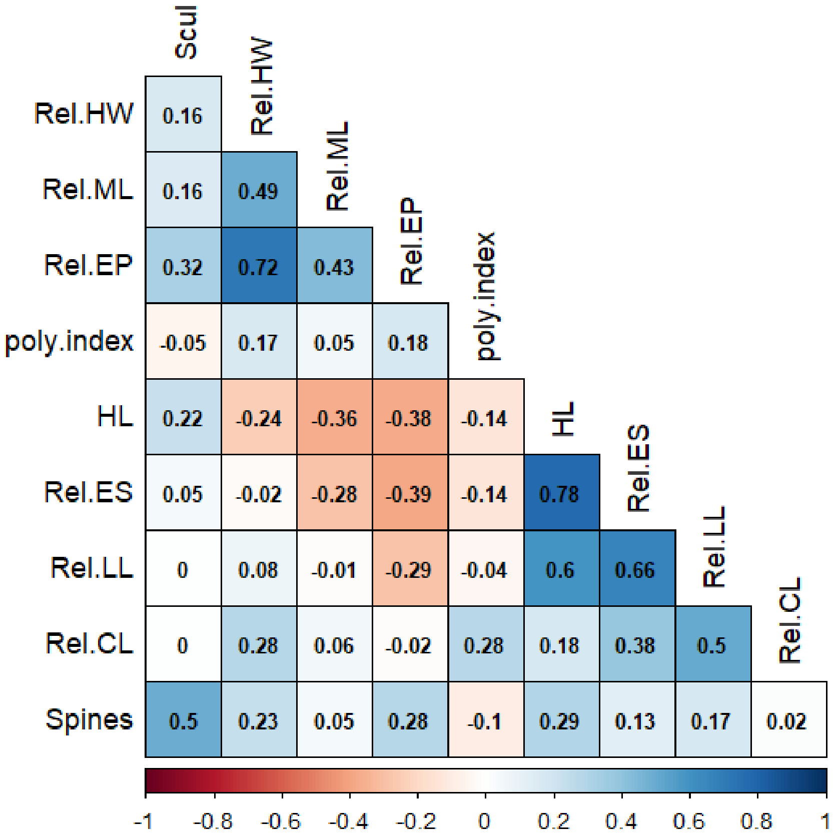


**Fig. S1.** Correlations between all traits used in this study. Given are spearman-rank coefficients, where more negative correlations are red while more positive correlations are blue. Trait abbreviations: Rel.HW = relative head width; Rel.ML = relative mandible length; Rel. EP: relative eye position; poly.index = polymorphisms index; HL = head length; Rel. ES = relative eye size; Rel. LL = relative leg length; Rel.CL = relative clypeus length.

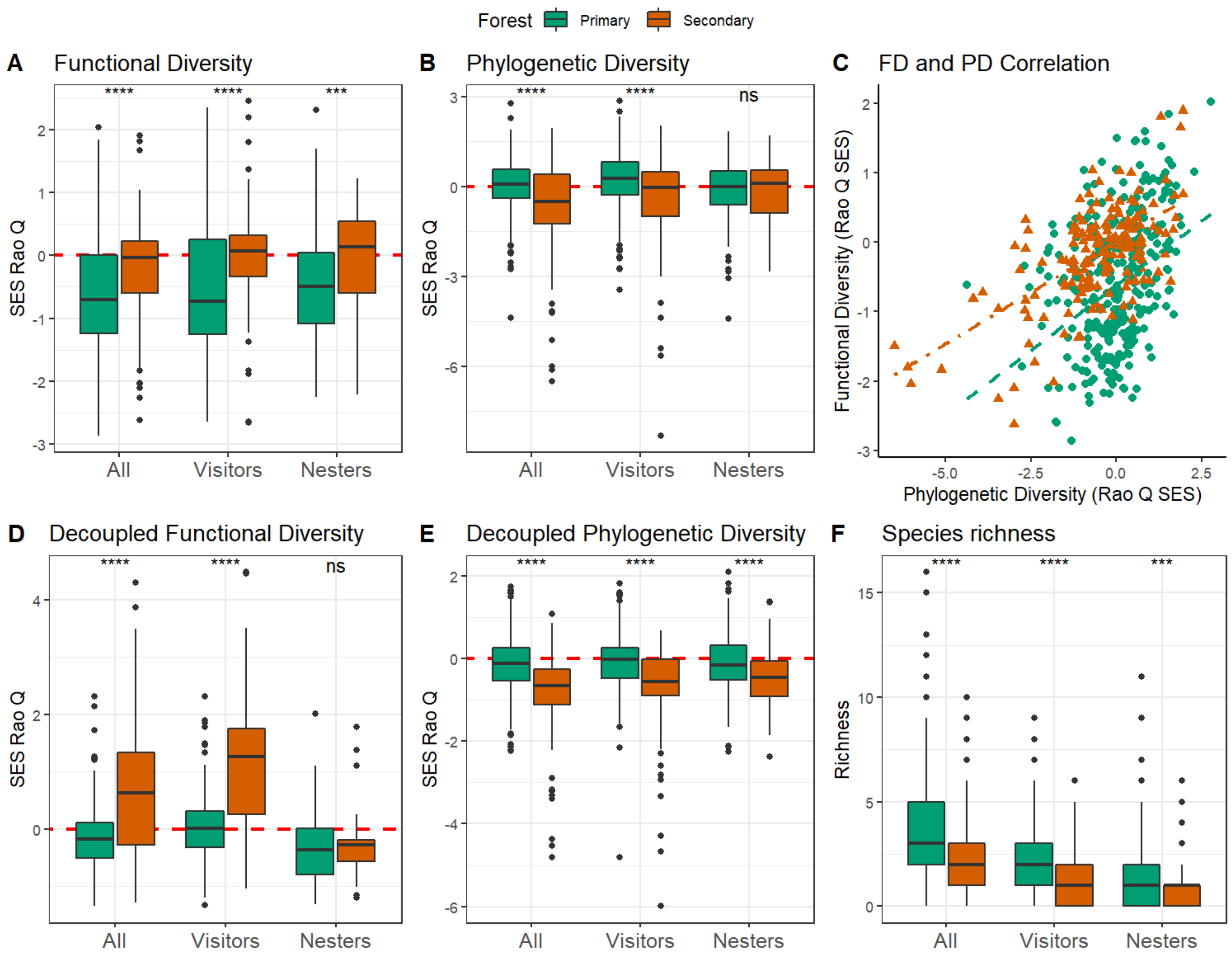


**Figure S2.** Arboreal ant functional diversity (A), phylogenetic diversity (B), the correlation between functional and phylogenetic diversity (C), the decoupled functional (D) and phylogenetic diversity (E) and the species richness (F) on tree scale after excluding all non-native species. Box-plots show median values per a tree with 25–75% quartiles and with whiskers representing 1.5 interquartile ranges for all species combined (All), for foraging species not nesting in a focal tree (Visitors) and nesting species (Nesters). Both forest types are shown in different colours (primary forest – green; secondary forest – orange) or symbols (in C: triangles denote primary forest, circles secondary forest). The values for individual ant communities are compared between the two forest types with a Kruskall-Wallis test, where stars indicate statistically significant differences (*** p < 0.001, ** p < 0.01, * p 0.05, ns p > 0.05). Standardised effect sizes (i.e., SES of Rao Q) are compared in A, B, D, and E where the dash horizontal line at 0 indicates random communities, while positive SES values indicate community overdispersion and negative values clustering. For mean observed and SES values and for tests of the statistical significance of the SES against null distribution, see Table 1.
